# From neuron to muscle to movement: a complete biomechanical model of *Hydra* contractile behaviors

**DOI:** 10.1101/2020.12.14.422784

**Authors:** Hengji Wang, Joshua Swore, Shashank Sharma, John Szymanski, Rafael Yuste, Thomas Daniel, Michael Regnier, Martha Bosma, Adrienne L. Fairhall

**Affiliations:** Department of Physics, University of Washington, Seattle, WA 98195; Department of Biology, University of Washington, Seattle, WA 98195; Department of Physiology and Biophysics, University of Washington, Seattle, WA 98195; NeuroTechnology Center, Department of Biological Sciences, Columbia University, New York 10027; Marine Biological Laboratory, Woods Hole, MA 02543; Department of Bioengineering, University of Washington, Seattle, WA 98195; Computational Neuroscience Center, University of Washington, Seattle, WA 98195

## Abstract

How does neural activity drive muscles to produce behavior? The recent development of genetic lines in *Hydra* that allow complete calcium imaging of both neuronal and muscle activity, as well as systematic machine learning quantification of behaviors, makes this small Cnidarian an ideal model system to understand and model the complete transformation from neural firing to body movements. As a first step to achieve this, we have built a biomechanical model of *Hydra*, incorporating its neuronal activity, muscle activity and body column biomechanics, incorporating its fluid-filled hydrostatic skeleton. Our model is based on experimental measurements of neuronal and muscle activity, and assumes gap junctional coupling among muscle cells and calcium-dependent force generation y muscles. With these assumptions, we can robustly reproduce a basic set of *Hydra’s* behaviors. We can further explain puzzling experimental observations, including the dual kinetics observed in muscle activation and the different engagement of ecto- and endodermal muscle in different behaviors. This work delineates the spatiotemporal control space of *Hydra* movement and can serve as a template for future efforts to systematically decipher the transformations in the neural basis of behavior.

## Introduction

To generate behaviors, neural activity is transformed through the biomechanics of the body. While there have been exciting developments of predictive models that incorporate musculoskeletal dynamics in humans and an array of vertebrate and invertebrate animals ***(Daniel, 1995; Rajagopal et al., 2016; Delp and Loan, 2000; Ting and Chiel, 2017)***, many are hampered by the inability to observe the system in its entirety, with simultaneous spatial and temporal patterns of neural activity, muscular activation and whole animal behavior. Small model systems pose the opportunity to develop relatively complete models of the transformations from neural activity to behavior, taking into account the biomechanics of the body ***(Kim et al., 2019; Pallasdies et al., 2019; Kim and Shlizerman, 2020)***. As an active medium, however, muscle tissue has its own dynamics that can contribute significantly to this transformation. The small freshwater *Cnidarian Hydra* offers a unique and unprecedented inroad into this problem. *Hydra* has a simple body structure; furthermore recently developed genetic lines now allow the direct imaging of calcium-based activity in both neurons ***(Dupre and Yuste, 2017)*** and muscle cells ***(Szymanski and Yuste, 2019)*** during behavior. This system thus presents an opportunity to account for the transformation from neural firing to behavior by modeling both body and active muscle dynamics, which determine how neural activation mediates behavioral outputs.

### Anatomy and behaviors of *Hydra*

*Hydra* has a relatively simple anatomy. Its fluid-filled body column consists of two body layers, the ectodermal and endodermal epithelia, separated and supported by an acellular gelatinous layer consisting of mesoglea, ***Figure 1***. The epithelial layers consist of a sheet of epitheliomuscular cells, innervated by separate nerve nets. The ectodermal and endodermal epitheliomuscular cells respectively produce longitudinal and circumferentially oriented contractions ***(Leclère and Röttinger, 2017)***. These layers, together with the enclosed fluid, form a hydrostatic skeleton in which the force of muscle contraction is transmitted throughout the body column by internal pressure ***(Kier, 2012)***.

**Figure 1.**
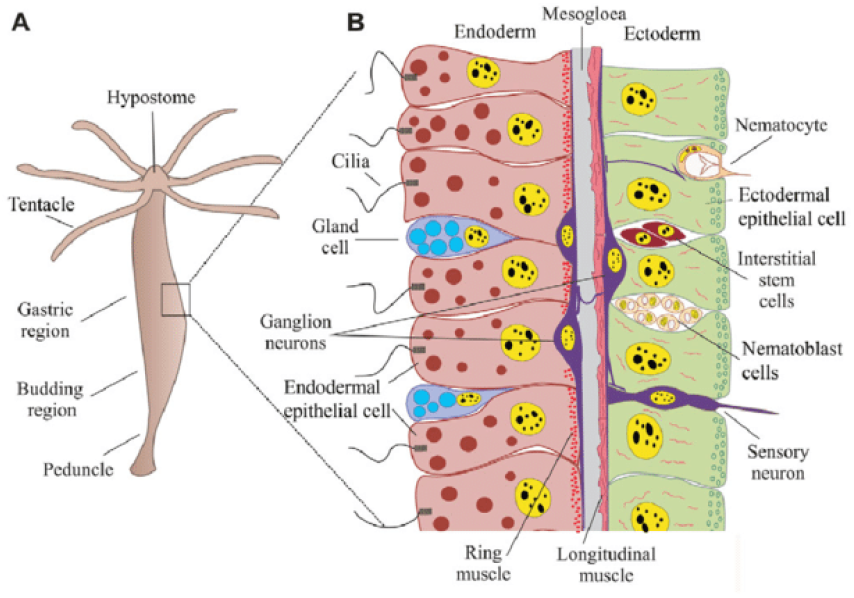
Anatomy of *Hydra **(Technau and Steele, 2011)***.

Movement in *Hydra* is controlled by a diffuse nerve net. *Hydra* has perhaps the first and simplest nervous system ***(Fujisawa and Hayakawa, 2012; Klimovich and Bosch, 2018)***; while its uncoupled ectodermal and endodermal nerve nets ***(Takaku et al., 2014)*** lack a centralized “brain” or ganglia, their firing activity underlies a rich repertoire of behaviors. These include contraction bursts, active elongation, nodding, bending and two forms of locomotion ***(Trembley, 1744; Rushforth et al., 1963; Passano and McCullough, 1964, 1965; Szymanski and Yuste, 2019; Han et al., 2018)***. However, how neural activity drives behavior is not understood. Early extracellular recordings identified several distinct electrical events in *Hydra:* contraction bursts (CB), rhythmic potentials (RP) and pre-locomotion bursts (PLB) ***(Passano and McCullough, 1962, 1963, 1964, 1965)***. Recent work has clearly identified three separate functional nerve subnetworks responsible for these electrical events ***(Dupre and Yuste, 2017)***. However, only one of these, the contraction burst (CB) network, is directly correlated with a motor output, namely whole-body contraction ***(Dupre and Yuste, 2017)***. Aside from that, the precise association of neural activity with behavior has not yet been mapped out.

### Neural control of behavior in *Hydra*

At what length scale and with what precision does the firing of nerve cells influence movement? These factors depend on how activation is conveyed through muscles, and how the resulting network of muscular contractions interacts with the biomechanics of the body. Due to gap junctional coupling ***(Westfall, 1973)***, the epitheliomuscular network is able to propagate excitation ***(Josephson, 1967; Josephson and Macklin, 1969; Anderson, 1980)*** even when *Hydra’s* nerve cells have been removed ***(Campbell et al., 1976; Lepault et al., 1980)***. Several studies have suggested that the contraction pulses can be conducted by the epithelium in Hydra ***Josephson, 1967; Josephson and Macklin, 1969; Kass-Simon, 1970, 1972; Rushforth, 1971)***; conduction in nerve-free epithelia has also been demonstrated in other hydrozoans such as Siphonophores, *Sarsia* and *Euphysa **(Mackie, 1965; Mackie and Passano, 1968)***. By imaging calcium signals in the endo- and ectodermal epithelial layers, ***Szymanski and Yuste (2019)*** reported two distinct forms of muscle layer activation: a rapid global activation that drives whole-body contraction, and slow waves of local activation that correlate with bending and possibly also elongation. This suggests that the dynamics of the muscle layer itself form an important and nontrivial component of the transformation from nerve firing to behavior.

Here, we construct a model of *Hydra* that includes a sufficient biophysical and biomechanical level of detail to simulate the complete transformation from neural activity to muscle activity to behavior. We aimed to address the following specific questions: (1) What are the mechanisms that support the observed dual timescales of muscle activation, and how does *Hydra* use these different dynamics in behavior? (2) During contraction bursts, although only neurons in the ectodermal nerve net fire ***(Dupre and Yuste, 2017)***, both muscle layers are activated ***(Szymanski and Yuste, 2019)***, and thus work against one another. What explains this joint activation and how can body contraction be achieved with opposing muscle drive? (3) Can we quantitatively reproduce basic behaviors ***(Han et al., 2018)***, including contraction, elongation and bending ?

To answer these questions, we implemented a multi-layered model, ***Figure 2***, transforming neural activity to movement. Our models are constrained both by observations from calcium imaging and by the use of physiologically plausible mechanisms consistent with the recently developed RNAseq database ***(Siebert et al., 2019)***. To model calcium dynamics in the epitheliomuscular cell network, we postulated the coexistence of a fast cellular electrically mediated pathway and a slow IP_3_-driven pathway. We assume that these activation signals are transmitted through the epithelial layers by gap junctions ***(Koenigsberger et al., 2004; Höfer et al., 2002; Loppini et al., 2015)***. These two mechanisms permit the coexistence of the fast electrically driven contractions as well as slow waves responsible for bending; the model predicts that these dynamics are triggered by distinct signals. We show that an intermediate level of gap junctional coupling between the ecto- and the endodermal epithelium can share contraction activation between the two muscle layers, but isolate slow wave activity to the ectoderm, consistent with observation. We next converted calcium dynamics to force generation. In order to account for *Hydra’s* movement dynamics, it was necessary to hypothesize that the relationship between calcium and force has more persistent dynamics in the endoderm than in the ectoderm, essentially suggesting slower relaxation times for endodermal muscles. Finally, we converted the simulated epithelial calcium dynamics to strain, which provided an active force input into a biomechanical model of the fluid-filled hydrostat. To our knowledge, such an actuated biological hydrostat has not been previously modeled. We use this model to show that the simulated muscle activation, when driven by neural activity inferred from calcium imaging, can account quantitatively for the measured behaviors of contraction, elongation and bending.

**Figure 2.**
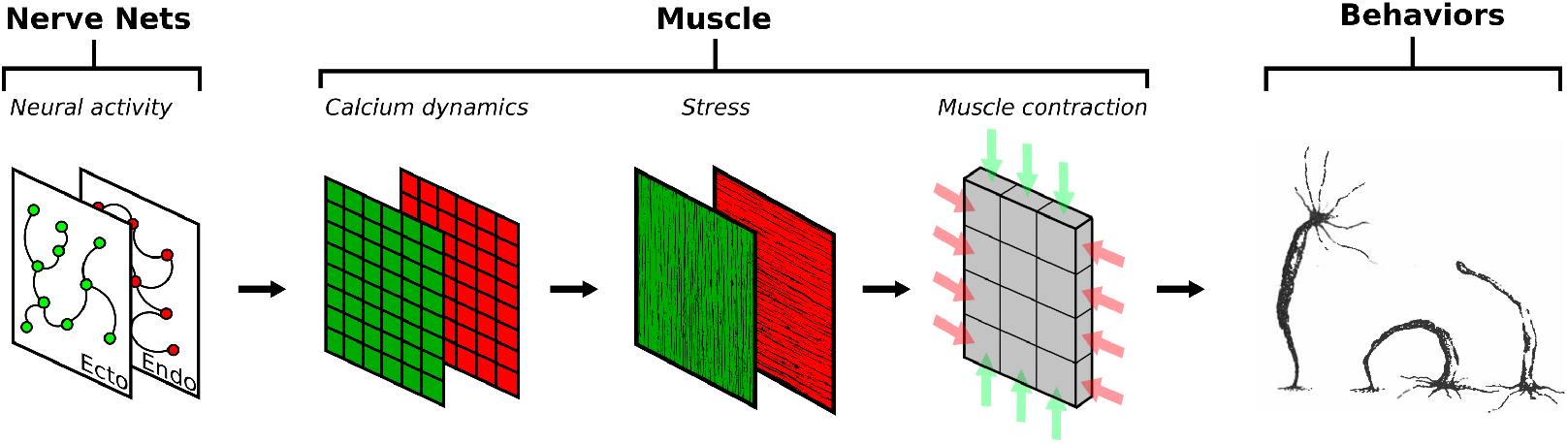
Framework of our project. Neural activity patterns trigger calcium dynamics in the muscle layers, which are transformed into contractile forces in the longitudinal (ectoderm; green) and circumferential (endoderm; red) directions, here indicated by the direction of fibres in the muscle layers. This provides the active force to drive a viscoelastic biomechanical model of the Hydra body column, simulating behaviors.

## Methods

Here we describe the development of our model in a sequence of steps, ***Figure 2***. We start from a biophysical model of calcium dynamics in a single epitheliomuscular cell. We then connect single cells into sheets via gap junctional coupling, and couple the two layers with sparsely distributed gap junctions. We convert calcium to strain using the Hai-Murphy model, and also convert it to fluorescence for comparison with imaging. Finally, we implement a biomechanical model of the *Hydra* body column and actuate it with active forces derived from the epithelial model. Finally, we describe data analysis methods used to extract information from *Hydra* calcium imaging for quantitative comparison with modeling results.

### Single cell modeling

*Hydra’s* main muscle cell type is the epitheliomuscular cell ***(Leclère and Röttinger, 2017)***, which has a similar structure to smooth muscle ***(Balachander et al., 2015)*** with no striation ***(Johnson et al., 2020)***. The ectodermal epitheliomuscular cells of *Hydra* display longitudinally oriented processes called myonemes which effect body contraction ***(Leclère and Röttinger, 2017)***. The myonemes of the endodermal epithelium are oriented circumferentially ***Figure 2***, and their contraction leads to elongation via conservation of volume of the interior fluid and corresponding hydrostatic pressure (the hydrostat mechanism).

Muscle contraction is controlled by calcium. Smooth muscle has two possible sources of calcium: (i) IP_3_-induced Ca^2+^ release from the endoplasmic reticulum/sarcoplasmic reticulum (ER/SR) calcium store, and (ii) Ca^2+^ influx from the extracellular space through L-type/T-type calcium channels ***(Horowitz et al., 1996; Berridge, 2008; Hill-Eubanks et al., 2011; Kuo and Ehrlich, 2015)***. We will refer to these two calcium signaling pathways as the “slow” and “fast” pathways respectively, based on their typical time scales. Both IP_3_-related calcium release ***Johnson et al., 2020)*** and electrical excitability ***(Holman and Anderson, 1991)*** are observed in epitheliomuscular cells.

Models of smooth muscle frequently treat only one of these pathways. Models of calcium signaling in non-excitable cells may consider only the slow pathway, ignoring membrane ion channels ***(De Young and Keizer, 1992; Li and Rinzel, 1994; Schuster et al., 2002; Handy et al., 2017)***, while others consider only the fast pathway, neglecting the dynamics of the internal calcium stores;examples include models for uterine smooth muscle ***(Rihana et al., 2009; Tong et al., 2011; Cochran and Gao,2015; Yochum et al.,2016; Testrow et al.,2018)***, gastric smooth muscle ***(Corrias and Buist, 2007)***, urinary bladder smooth muscle ***(Mahapatra et al., 2018)*** and pancreatic *β*-cells ***(Fridlyand et al., 2009)***. While some models integrate both pathways by incorporating both influx through ion channels and Ca^2+^ release from stores ***(Imtiaz et al., 2006; Kusters et al., 2005; Koenigsberger et al., 2004)***, modeling the two pathways separately and simulating calcium dynamics at different time scales is rare ***(Fletcher and Li, 2009; Halidi et al., 2011)***. However, since calcium imaging in *Hydra* clearly reveals dynamics with different time scales (short-lasting and fast-propagated calcium transients in CB;long-lasting, slow calcium waves in bending and nodding ***(Szymanski and Yuste, 2019)***, we hypothesize that both the slow and fast pathways coexist in *Hydra* epitheliomuscular cells.

#### Fast Pathway

The fast pathway of our model is initiated by the binding of neuropeptides on ionotropic receptors, which activate ligand-gated ion channels (LGIC) and depolarize the cellular membrane ***(Gründer and Assmann, 2015)***. The elevated membrane potential activates calcium channels (L-type/T-type), triggering a large influx of Ca^2+^ from the extracellular space and invoking a further depolarization. Meanwhile, the high membrane potential inactivates calcium channels and activates potassium channels, resulting in membrane repolarization ***(Hill-Eubanks et al., 2011)***. Ca^2+^ is extruded to the extracellular space through plasma membrane Ca^2+^ ATPase (PMCA) and is recycled to the ER through sarco/endoplasmic reticulum Ca^2+^-ATPase (SERCA) ***(Fujisawa and Hayakawa, 2012; Dupont et al., 2016)***.

#### Slow Pathway

In the slow pathway, neuropeptides bind a G protein-coupled receptor (GPCR) and activate a G protein, which activates phospholipase C (PLC) and hydrolyzes phosphatidylinositol bisphosphate (PIP2) into inositol 1,4,5-trisphosphate (IP_3_), which plays a role of the second messenger for calcium signaling. IP_3_ can bind to IP_3_ receptors (IPR) on the endoplasmic reticulum (ER) membrane and thus cause the release of Ca^2+^ from the ER. In this pathway, Ca^2+^ is also extruded through PMCA and recycled through SERCA.

We combined these mechanisms in an intracellular calcium signaling model ***Equation 1 (Figure 3A)***, where the dynamical variables are cytosolic Ca^2+^ concentration *(C)*, ER Ca^2+^ concentration *(S)*, cytosolic IP_3_ concentration *(P)* and membrane voltage *(V)*. The terms are defined in ***Table 1***.

**Figure 3.**
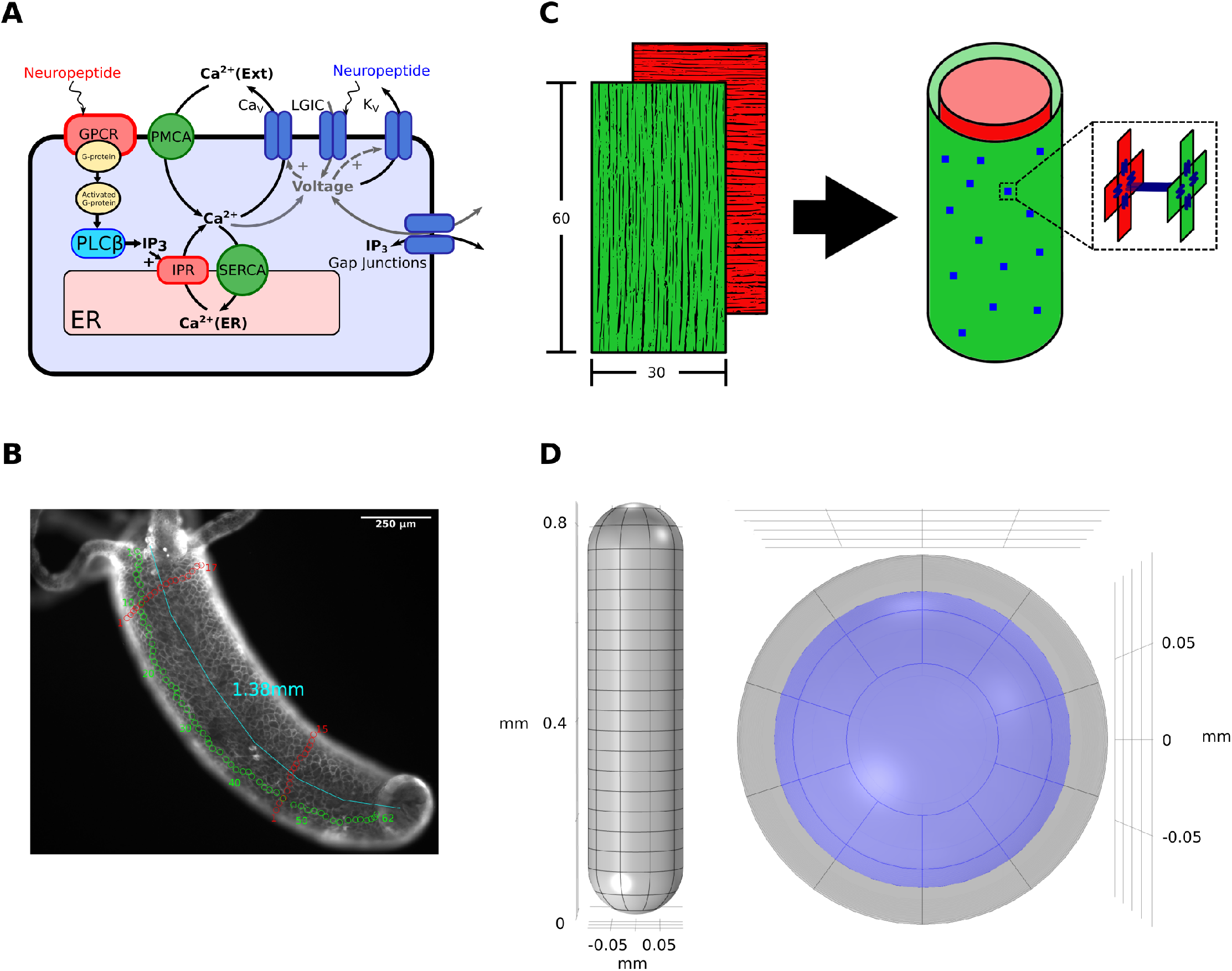
Modeling details. *(A)* Intracellular calcium signaling model including both pathways. Fast and slow pathways are separately triggered by neuropeptides (red and blue, respectively). *(B)* Cell counting and length measurement in a small *Hydra.* Green circles are the counts in longitudinal direction and red colors are the counts in circumferential direction. *(C)* The layout of muscle networks. Green and red represent the ectodermal and endodermal layers respectively, in which the myonemes are respectively aligned in longitudinal and circumferential directions, as represented by the fibre directions. Blue dots represent cross-layer gap junctions. *(D)* Geometry of the biomechanical model from a side view (left) and top view (right). Gray represents the muscle shell domain, while the enclosed fluid is shown as purple.

**Table 1.** Terms involved in ***Equation 1***

| Term | Expression | Description |
| --- | --- | --- |
| $J_{\text{IPR}}$ | $k_{\text{IPR}} \frac{C^2 I^2 R}{(C^2 + K_a^2)(I^2 + K_{\text{IP}}^2)} (S - C)$ | Ca<sup>2+</sup> flux through IPR |
| $J_{\text{leak}}$ | $k_{\text{leak}} (S - C)$ | Ca<sup>2+</sup> leak out of ER |
| $J_{\text{SERCA}}$ | $k_{\text{SERCA}} C$ | Ca<sup>2+</sup> flux through SERCA pump |
| $J_{\text{PMCA}}$ | $k_{\text{PMCA}} C$ | Ca<sup>2+</sup> flux through PMCA pump |
| $J_{\text{in}}$ | $v_{\text{in}} + v_r \frac{P^2}{K_r^2 + P^2}$ | Ca<sup>2+</sup> entry fluxes |
| $v_{\text{PLC}\beta}$ | $v_{\beta}$ | Rate of PLCβ |
| $v_{\text{deg}}$ | $k_{\text{deg}} P$ | Rate of IP<sub>3</sub> degradation |
| $I_{\text{Ca}}$ | $g_{\text{Ca}} m^2 h (V - E_{\text{Ca}})$ | Current through Ca<sup>2+</sup> channels |
| $I_{\text{K}}$ | $g_{\text{K}} n^4 (V - E_{\text{K}})$ | Current through K<sup>+</sup> channels |
| $I_{\text{L}}$ | $g_{\text{L}} (V - E_{\text{L}})$ | Current through leak channels |
| $I_{\text{stim}}$ | 0 (resting) or 0.01 (stimulated) mA/cm<sup>2</sup> | Current through ligand-gated ion channels |

Source: Expressions of $J_{\text{IPR}}$ , $J_{\text{leak}}$ , $J_{\text{SERCA}}$ , $J_{\text{PMCA}}$ , $J_{\text{in}}$ , $v_{\text{PLC}\beta}$ , $v_{\text{deg}}$ are from *Höfer et al. (2002)*; $I_{\text{Ca}}$ and $I_{\text{K}}$ is modified from *Diderichsen and Göpel (2006)*.

In our model, these two pathways are independent: when calcium is elevated in the slow pathway, it will not induce calcium influx in the fast pathway, and vice versa. For the slow pathway, the opening of IPR requires a high concentration of IP_3_, so calcium elevation caused by influx from outside cannot trigger a calcium release from ER; for the fast pathway, calcium channels are only dependent on the membrane potential, so the slow pathway can not trigger calcium influxes. One potential coupling mechanism would be calcium-activated chloride channels; however, gene-query of *Hydra’s* genome does not support the presence of these channels. We further assume that the two pathways can be independently triggered by different types of neuropeptides, based on previous identification of distinct *Hydra* peptide functions ***(Fujisawa and Hayakawa, 2012)***.

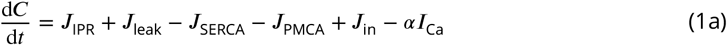

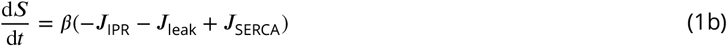

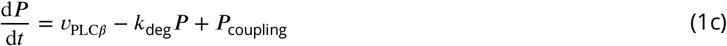

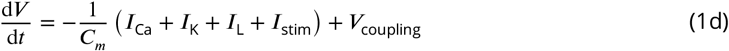

The model also includes the inactivation rate of IPR (*R*) in *J*_IPR_, the activation *(m)* and inactivation *(h)* gating variables in *I_Ca_*, the activation (*n)* gating variable in *I_K_*. The dynamics of *R* are:

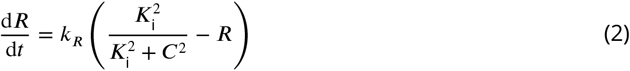

The dynamics of any gating variable *g* ∈ *{m,h,n}* can be represented as

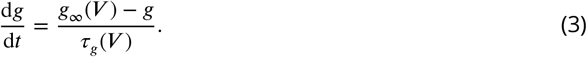

Expressions for the steady-state values *g_∞_(V)* (unitless) and the time constants *τ_g_(V)* ([s]) are shown in ***Equation 4***. The values and descriptions of all corresponding parameters above can be found in ***Table 2***.

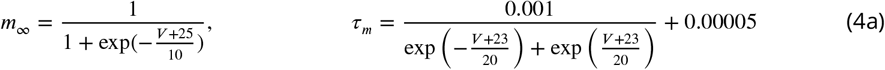

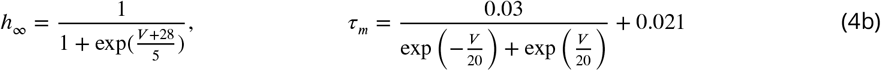

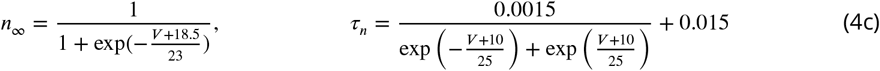

**Table 2.** Parameters used in modeling calcium dynamics

| Parameter | Description | Value | Unit |
| --- | --- | --- | --- |
| $k_{\text{IPR}}$ | Rate constant of $\text{Ca}^{2+}$ release through IPR | 0.08 | $\text{s}^{-1}$ |
| $k_R$ | Rate constant of IPR inactivation | 4 | $\text{s}^{-1}$ |
| $K_a$ | Half-saturation constant for $\text{Ca}^{2+}$ activation of IPR | 0.2 | $\mu\text{M}$ |
| $K_i$ | Half-saturation constant for $\text{Ca}^{2+}$ inhibition of IPR | 0.2 | $\mu\text{M}$ |
| $K_{\text{IP}}$ | Half-saturation constant for $\text{IP}_3$ activation of IPR | 0.3 | $\mu\text{M}$ |
| $k_{\text{leak}}$ | Rate constant of $\text{Ca}^{2+}$ leak from ER | 0.0004 | $\text{s}^{-1}$ |
| $k_{\text{SERCA}}$ | Rate constant of SERCA pump | 0.3 | $\text{s}^{-1}$ |
| $k_{\text{PMCA}}$ | Rate constant of PMCA pump | 0.8 | $\text{s}^{-1}$ |
| $v_{\text{in}}$ | Rate of $\text{Ca}^{2+}$ leak across the plasma membrane | 0.025 | $\mu\text{M/s}$ |
| $v_r$ | Maximal rate of activation-dependent $\text{Ca}^{2+}$ influx | 0.2 | $\mu\text{M/s}$ |
| $K_r$ | Half-saturation constant for activation-dependent $\text{Ca}^{2+}$ entry | 1 | $\mu\text{M}$ |
| $v_\beta$ | Rate of $\text{PLC}\beta$ | 0.002 - 1.0 | $\mu\text{M/s}$ |
| $k_{\text{deg}}$ | Rate constant of $\text{IP}_3$ degradation | 0.05 | $\text{s}^{-1}$ |
| $\alpha$ | Current conversion factor | 5182.13 | $\mu\text{M}\cdot\text{cm}^2/\text{mC}$ |
| $\beta$ | Ratio of the volumes of cytoplasm and ER | 20 | 1 |
| $g_{\text{Ca}}$ | Maximal conductance of $I_{\text{Ca}}$ | 0.0005 | $\text{S/cm}^2$ |
| $E_{\text{Ca}}$ | Reversal potential of $I_{\text{Ca}}$ | 51 | mV |
| $g_{\text{K}}$ | Maximal conductance of $I_{\text{K}}$ | 0.0025 | $\text{S/cm}^2$ |
| $E_{\text{K}}$ | Reversal potential of $I_{\text{K}}$ | -75 | mV |
| $g_{\text{L}}$ | Conductance of $I_{\text{L}}$ | 0.000036 | $\text{S/cm}^2$ |
| $E_{\text{L}}$ | Reversal potential of $I_{\text{L}}$ | -55 | mV |
| $C_m$ | Membrane capacitance per unit area | 1 | $\mu\text{F/cm}^2$ |

#### Neuromuscular coupling

The mechanisms of communication between the nerve nets and epitheliomuscular layers are not completely known. Neuromuscular synaptic junctions have been observed using electron microscopy ***(Westfall, 1973)***. A number of signalling molecules appear to mediate the interaction between nerve activity and muscle excitation. The “Hydra Peptide Project” identified distinct functions of different neuropeptides ***(Fujisawa and Hayakawa, 2012)***; Hym-153 (FRamide-2) is believed to directly act on body contraction, while Hym-65 (FRamide 1) and Hym-248 are thought to act on body elongation.

In our model, we simulated these different interactions by triggering the fast pathway with a step of input current *I*_stim_ from 0 to 0.01 mA/cm^2^ for 10ms, to simulate the influx of positive ions through the temporary opening of ligand-gated ion channels triggered by neuropeptide stimulation; we triggered the slow pathway by elevating *υ_β_* for 4s, which represents the process from the binding of neuropeptides on GPCR until the activation of PLC*β*.

### Muscle network modeling

From the single cell model, we then proceeded to build a multicellular model for the muscle sheets of *Hydra*, in order to explain the activation patterns recorded using calcium imaging ***(Szymanski and Yuste, 2019)***. We assume that the epitheliomuscular cells are connected by gap junctions; these have been observed in EM studies ***(Takaku et al., 2014)*** between cells in the same layer, and also penetrating the mesoglea to connect the ectoderm and endoderm ***(Hand and Gobel, 1972; Wood and Kuda, 1980; Takaku et al., 2014)***. Further support for the involvement of gap junctions comes from single-cell RNA sequence analysis in *Hydra*, which shows multiple innexin types in both epithelial cells and neurons ***(Siebert et al., 2019)***. In most systems, nerve stimulation alone does not activate the majority of smooth muscle cells; rather, activation is propagated via intercellular communications among muscle cells ***(Anderson, 1980; Christ et al., 1996)***. Gap junctions propagate signals by (1) allowing the diffusion of Ca^2+^ as well as second messengers like IP_3_; (2) conducting electrical signals ***(Anderson, 1980; Huizinga et al., 1992; Christ et al., 1996)***.

We hypothesize that the two different observed forms of wave propagation in *Hydra* (slow waves and fast calcium synchronization) both occur through different epitheliar activation patterns. For slow waves, the propagation of IP_3_, but not Ca^2+^ ***(Jafri and Keizer, 1994)***, through gap junctions is believed to trigger intercellular calcium waves (ICW) ***(Leybaert and Sanderson, 2012)***, which is supported by many models ***(Sneyd et al., 1995; Dupont et al., 2000; Höfer et al., 2001,2002; Goldberg et al., 2010)***. Calcium synchronization (the fast wave) has generally been modelled through the electrical conduction property of gap junctions ***(Imtiaz et al., 2006; Kusters et al., 2008; Koenigsberger et al., 2010)***.

To simulate calcium signaling in the whole-body muscle sheets, we constructed ectodermal and endodermal networks of *Hydra* epitheliomuscular cells. The number of muscle cells of *Hydra* varies considerably with body size ***(Tzouanas et al., 2019)***. For a representative *Hydra* of length of 1.38 mm, ***Figure 3***B, we counted 62 cells longitudinally and 30 (15×2)-34 (17×2) cells circumferentially, depending on the longitudinal location. We approximated the body column as a cylinder composed of 30×60 square cells, of which the side length is 30 *μ*m; the lateral sheet edges are connected and the topmost and bottommost cell rows are taken to be isolated from the environment (***Figure 3***C). Cells within a layer are connected to their neighbours via gap junctions.

Each cell is treated as a compartment. To model the role of gap junctions in propagating electrical signals and diffusing IP_3_, neighbouring cells within a layer and the cells at the same location in the endoderm and ectoderm are connected by coupling terms 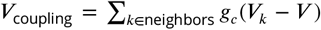 and 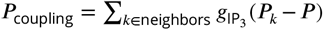 (incorporated in ***Equation 1***). While all neighboring cells within a layer are coupled, cells with the same indices in the two layers are connected probabilistically, with a defined connectivity ratio. The gap junctional conductances *g_c_* and *g*_IP_3__ are inhomogeneous in different directions and different layers; the specific values can be found in ***Table 2***.

### Force generation

We next model how calcium dynamics drive muscle contraction. In smooth muscle, a rise of [Ca^2+^]_*i*_ (intracellular calcium concentration) leads to a rise of calmodulin, leading to increased activation of MLCK (myosin light-chain kinase), phosphorylation of myosin, and thus contraction ***(Dupont et al., 2016)***. The Hai-Murphy model of smooth muscle ***(Hai and Murphy, 1988)*** uses four forms of the crossbridge to simulate the force-production process, where Ca^2+^ plays a role in MLCK activation. The Hai-Murphy model includes the “latch-state” of the crossbridge, which allows the maintenance of steady-state stress of muscle even if the Ca^2+^ concentration has decreased. Depending on parameters, the model can provide either fast-acting phasic behavior as for “fast” myosin isoforms, or tonic contraction via the latch-bridge mechanism, as for “slow” isoforms. This mechanism is energetically efficient but at the cost of a reduced rate of muscle shortening ***(Han et al., 2006)***. Modified Hai-Murphy models have been used to simulate contractions in arteriole ***(Wang et al., 2008)*** and uterine ***(Maggio et al., 2012; Yochum et al., 2016)*** smooth muscle.

We applied a modified version of the Hai-Murphy model, following ***Maggio et al. (2012)*** and ***Yochum et al. (2016)***, to transform [Ca^2+^]_*i*_ into active stress. ***Equation 5*** governs the dynamics of the four states of the crossbridge, unattached and unphosphorylated *(M)*, unattached and phosphorylated *(M_p_)*, attached and phosphorylated *(AM_p_)*, attached and unphosphorylated *(AM)*, as follows:

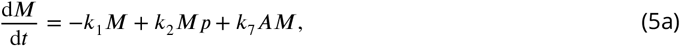

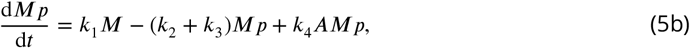

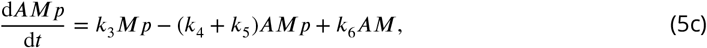

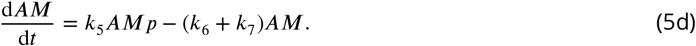

The final active stress *F_a_* is calculated as

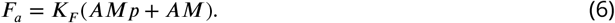

The values and descriptions of parameters in the modified Hai-Murphy model are shown in ***Table 3***. Contractions in ectoderm and endoderm have different durations: the active elongation phase of *Hydra* can last for hundreds of seconds despite a decay of the calcium concentration, indicating a maintained tonic contraction in the endoderm, while each contraction pulse initiates and decays rapidly. This is reminiscent of “slow” and “fast” myosin isoforms in ***Han et al. (2006)***, explained by the existence and detachment rate of the “latch-bridge” state, parametrized by *fc*_7_ in the Hai-Murphy model. We also set different values of 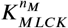 in the two layers, assuming that the [Ca^2+^]_*i*_ sensitivities of ectodermal and endodermal MLCK are different. We further tune additional parameters in the two muscle layers to fit the length change of the *Hydra* body during different behaviors (see Results). Overall, through these choices, endodermal contractions are more easily activated and maintained than those in the ectoderm.

**Table 3.** Parameters used in the modified Hai-Murphy model

| Parameter | Description | Value | Unit |
| --- | --- | --- | --- |
| $k_1$ | Rate of $M \rightarrow M_p$ | $\frac{C^{n_M}}{K_{MLCK}^{n_M} + C^{n_M}}$ | $s^{-1}$ |
| $k_2$ | Rate of $M_p \rightarrow M$ | 0.15 | $s^{-1}$ |
| $k_3$ | Rate of $A + M_p \rightarrow AM_p$ | 16 (ecto) / 0.4 (endo) | $s^{-1}$ |
| $k_4$ | Rate of $AM_p \rightarrow A + M_p$ | 4 (ecto) / 0.05 (endo) | $s^{-1}$ |
| $k_5$ | Rate of $AM_p \rightarrow AM$ | $k_2$ | $s^{-1}$ |
| $k_6$ | Rate of $AM \rightarrow AM_p$ | $k_1$ | $s^{-1}$ |
| $k_7$ | Rate of $AM \rightarrow A + M$ | 0.75 (ecto) / 0.015 (endo) | $s^{-1}$ |
| $K_{MLCK}^{n_M}$ | Half-saturation $[Ca^{2+}]_i$ for MLCK activation | 0.85 (ecto) / 0.15 (endo) | $\mu M$ |
| $n^M$ | Hill coefficient of $Ca^{2+}$ activation of MLCK | 4 | 1 |
| $K_F$ | Proportional coefficient of active stress generation | 3.3 (ecto) / 0.3 (endo) | $N/mm^2$ |

### Transformation of [Ca^2+^]_*i*_ to fluorescence

To better compare our simulations directly with results from calcium imaging, we convert calcium concentration into fluorescence. In *Hydra* muscle calcium imaging experiments, GCaMP6s and jRCaMP1b were used as indicators in the ectoderm and endoderm respectively ***(Szymanski and Yuste, 2019)***. We used the fluorescence sequential binding model (SBM) of ***Greenberg et al. (2018**) (**Equation 7)***, in which five dynamical variables *G_0_* - *G*_4_ represent the concentration of possible GCaMP6s binding states, with 0 to 4 ions bound:

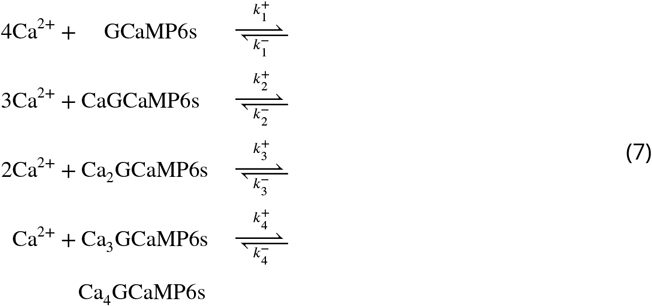

Following ***Greenberg et al. (2018)***, we use *r_j_*(*j* = 1,2,3,4) to represent the rates of transition from *G_j-1_* to *G_j_*:

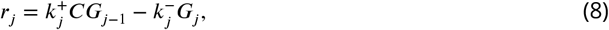

resulting in:

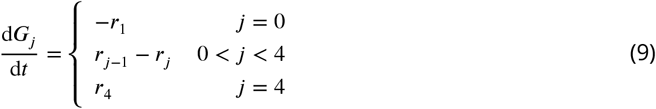

The fluorescence intensity *F* is computed as

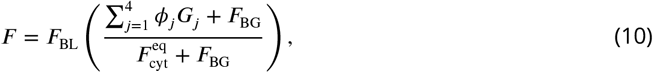

where the definitions and values of the parameters are shown in ***Table 4***.

**Table 4.** Parameters used in the SBM

| Parameter | Description | Value | Unit |
| --- | --- | --- | --- |
| $k_1^+$ | First GCaMP6s on-rate | 20 | $\mu\text{M}^{-1}\text{s}^{-1}$ |
| $k_1^-$ | First GCaMP6s off-rate | 0.1 | $\mu\text{M}^{-1}\text{s}^{-1}$ |
| $k_2^+$ | Second GCaMP6s on-rate | 16.9 | $\mu\text{M}^{-1}\text{s}^{-1}$ |
| $k_2^-$ | Second GCaMP6s off-rate | 20 | $\mu\text{M}^{-1}\text{s}^{-1}$ |
| $k_3^+$ | Third GCaMP6s on-rate | 1.1 | $\mu\text{M}^{-1}\text{s}^{-1}$ |
| $k_3^-$ | Third GCaMP6s off-rate | 5 | $\mu\text{M}^{-1}\text{s}^{-1}$ |
| $k_4^+$ | Fourth GCaMP6s on-rate | 1069 | $\mu\text{M}^{-1}\text{s}^{-1}$ |
| $k_4^-$ | Fourth GCaMP6s off-rate | 2 | $\mu\text{M}^{-1}\text{s}^{-1}$ |
| $\phi_1/\phi_0$ | CaGCaMP6s/GCaMP6s brightness ratio | 1.0 | 1 |
| $\phi_2/\phi_0$ | Ca<sub>2</sub>GCaMP6s/GCaMP6s brightness ratio | 1.0 | 1 |
| $\phi_3/\phi_0$ | Ca<sub>3</sub>GCaMP6s/GCaMP6s brightness ratio | 1.0 | 1 |
| $\phi_4/\phi_0$ | Ca<sub>4</sub>GCaMP6s/GCaMP6s brightness ratio | 81.0 | 1 |
| $F_{\text{BG}}/F_{\text{cyt}}^{\text{eq}}$ | Background/cytosolic fluorescence at rest | 2.5 | 1 |

### Biomechanical modeling

Our final step is to develop simulations of the biomechanics of the body. Models of hydrostatic skeletons are rare and have focused on muscular hydrostats, including leech ***(Wadepuhl and Beyn, 1989; Skierczynski et al., 1996)*** and octopus tentacles ***(Yekutieli et al., 2005)***. Related work has explored the neural control of swimming in jellyfish ***(Pallasdies et al., 2019)***; in this case the animal is an open membrane and the model treats the hydrodynamic interaction with the surrounding fluid. We constructed our model using the COMSOL Multiphysics^®^ 5.3 package.

We approximated the anatomy of *Hydra* with a simplified biomechanical model that contains two domains: the body shell and the enclosed fluid. The body shell, which represents the combination of the ectoderm and endoderm layers and the mesoglea, is composed of a half spherical shell at the hypostome and a half spherical shell at the peduncle, connected by a uniform body column cylinder shell (***Figure 3***D). In order to manipulate the biomechanical model at high resolution, we divided the body shell into 10 (radial) by 20 (longitudinal) elements. The body shell and enclosed fluid together form a hydrostatic skeleton.

To define the passive properties of *Hydra* body, we defined the body shell of our model as an incompressible hyperelastic material which follows a Neo-Hookean model ***(Rivlin, 1948)***. Hyperelastic materials exhibit a nonlinear stress-strain behavior and can respond elastically under very large strains ***(Bower, 2009)***. Muscle tissues are often well-described ***(Gras et al., 2012; Sarma et al., 2003; Chagnon et al., 2015)*** and modeled ***(Ansari et al., 2007; Gras et al., 2010; Tang et al., 2009)*** using hyperelastic properties. The passive biomechanical properties were mostly modeled based on Hill’s three element model ***(Hill, 1938)***. Since biological soft tissues have hyperplasticity or viscoelasticity ***(Martinek et al., 2008)***, we used hyperelastic material parameters to model the *Hydra* muscle shell and further incorporated viscoelasticity by including a Kelvin-Voigt model into the body shell material. For the enclosed fluid, we used the COMSOL simulation environment’s preset material “Water”, with a moving mesh. Parameters of the materials are shown in ***Table 5***.

**Table 5.** Parameters used in the biomechanical model

| Parameter | Value | Unit |
| --- | --- | --- |
| Top half sphere radius | 97.5 | $\mu\text{m}$ |
| Bottom half sphere radius | 97.5 | $\mu\text{m}$ |
| Body cylinder height | 650 | $\mu\text{m}$ |
| Thickness | 19.5 | $\mu\text{m}$ |
| Density of Enclosed Fluid | 1000 | $\text{kg}/\text{m}^3$ |
| Density of Muscle | 1200 | $\text{kg}/\text{m}^3$ |
| Lamé parameter $\mu$ of Muscle | 3.336 | kPa |
| Lamé parameter $\lambda$ of Muscle | 1.664 | GPa |
| Initial bulk modulus of muscle | 1.667 | GPa |
| Relaxation time of muscle | 70 | s |

While the COMSOL architecture as described handles the passive biomechanical properties, we applied time-varying active stresses generated by the output of the calcium signaling models; the ectodermal model drove longitudinal external stresses, while the endodermal output drove stress in the circular direction. We obtained the active stress from the calcium signaling model as described above, and averaged the stress of neighboring 9 cells, coarse-graining the original 30×60 matrix to fit the 10×20 dimensions of the biomechanical model. This stress pattern was applied to the corresponding elements of the biomechanical model using LiveLink^TM^ for MATLAB.

### Model constraints: gene-expression database

To validate choices of biophysical mechanisms including channels, receptors and pumps, we queried gene expression information for proposed components. We identified candidate genes by FASTA using the NCBI protein database and then used BLAST ***(Altschul et al., 1990)*** to search for these genes in the databases *Hydra* 2.0 genome, Augustus Gene Models and Juliano aepLRv2. The Broad *Hydra* Single-Cell Portal ***(Siebert et al., 2019)*** further allowed us to identify body regions with corresponding gene expression. We limited ourselves to mechanisms that were consistent with these data bases.

### Video analysis of muscle activity and behavior

#### Hydra cultures and imaging

All Hydra lines were maintained at 18°C and fed newly hatched *Artemia nauplii* two to three times per week. *Hydra* expressing the calcium indicator GCaMP6 in the ectoderm of the animal were used for imaging experiments. We used a modified imaging preparation from ***Dupre and Yuste (2017)***. All imaging took place under a ZEISS Axio Zoom.V16 equipped with Zeiss AxioCam 506 monochrome camera for fluorescent imaging, PlanNeoFluar Z 2.3X objective lens and a GFP fluorescent filter set. The imaging arena consisted of a microscope slide, 50 to 100 μm spacer and a cover slip. The use of the spacer allowed us to keep the animals in focus by preventing motion in the z direction while still allowing free motion in the *x* and *y* directions. Animals were recorded in the arena for 30-60 minutes at a sampling rate of 4 to 10 frames per second.

#### Video Analysis

Here we use image analysis to estimate integrated fluorescence in the neuronal and muscle GCAMP lines (in contrast to single neuron tracking ***(Lagache et al., 2020**))* as well as to accurately characterize *Hydra’s* body configuration. Acquired movies were processed using a combination of ImageJ ***(Schneider et al., 2012)***, the Icy Imaging software suite ***(De Chaumont et al., 2012)***, DeepLabCut ***(Mathis et al., 2018)*** and custom scripts, with a pipeline shown in ***Figure 4***. ImageJ was used to adjust the contrast from background noise which is essential to accurately extract contours of the *Hydra.* Noise was reduced using median filtering (despeckle plugin). Icy Imaging was then used to extract the contours of individual frames using the Active Contours plugin. We can then integrate fluorescence signals within the contour. We then used DeepLabCut to track 4 reasonably well-identified body locations: the center of the hypostome, the center of the peduncle, and the points of intersection of the left- and rightmost tentacles with the body column (the “armpits”). The tracked “armpits” from DeepLabCut were used to exclude the tentacles from the Icy contour. We then used the peduncle to bisect the contour, and proportionally segment the two contour halves. Connecting the midpoints of the segmentation points allowed us to extract the curved midline of the *Hydra* body in each frame.

**Figure 4.**
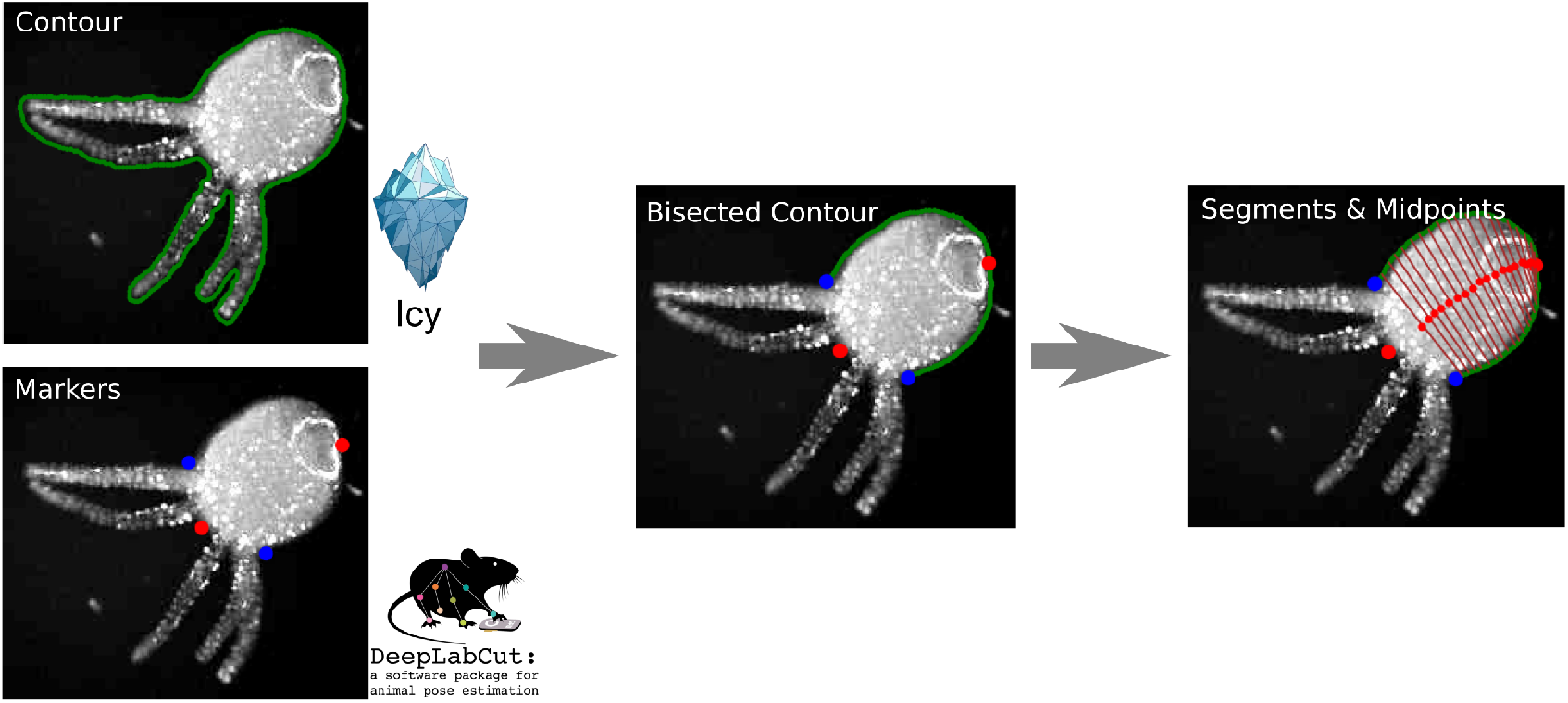
Pipeline of the video analysis.

## Results

We implemented the sequence of models as described in Methods and illustrated in ***Figure 2***.

### Single cell dynamics

We implemented the single cell model shown in ***Figure 3***A. This model has two pathways that drive calcium dynamics: a fast pathway by which calcium enters through voltage-gated ion channels, and a slow pathway, triggered by neuropeptide acting on a G-protein coupled receptor and mediated by IP_3_ (see Methods), releasing calcium from intracellular stores. We show these dynamics in ***Figure 5***, demonstrating the different timescales of the [Ca^2+^]_*i*_ transient in the two pathways.

**Figure 5.**
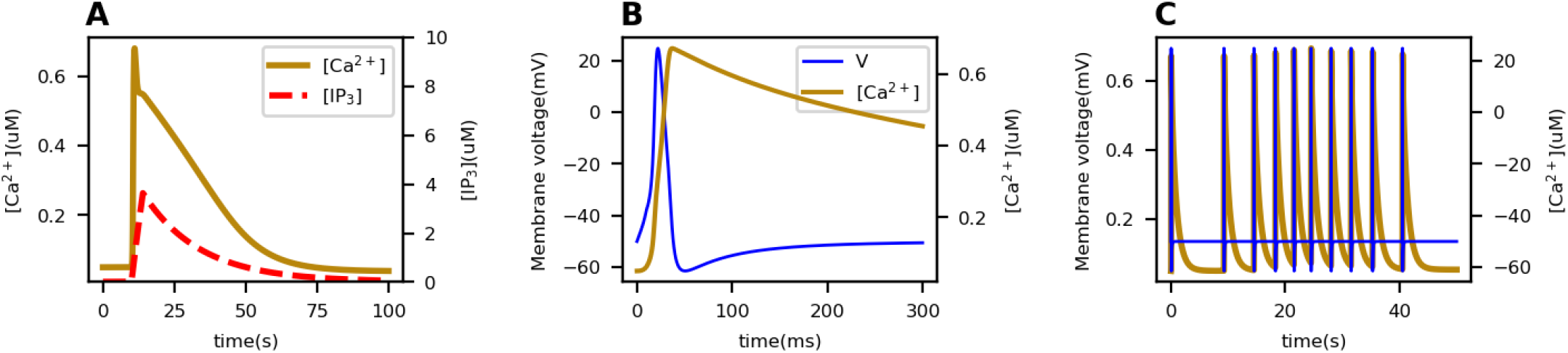
Dynamics of a single muscle cell in *Hydra. (A)* Concentrations of Ca^2+^ and IP_3_ when slow pathway is activated. *(B)* Membrane potential (blue) and Ca^2+^ concentration (brown) in a single spike when fast pathway is activated. *(C)* Membrane potential (blue) and Ca^2+^ concentration (brown) under multiple pulses of stimulation of fast pathway.

### Muscle network dynamics

We now show how the observed muscle layer dynamics can be successfully reproduced by our model.

#### Body column wave

To simulate body-column slow waves in our model, we hypothesize that their initiation arises from neuromuscular junctions at which neurons release a neuropeptide that triggers the slow pathway dynamics in the joint muscle cells. To initiate these waves, we randomly select a 2×2 region of cells in the sheet and stimulate their slow pathway calcium dynamics. The elevated IP_3_ in the stimulated cells diffuses to the neighboring cells and triggers the slow dynamics there, resulting in slow calcium waves propagating through the corresponding local domains. An example result of the simulation is shown in ***Figure 6***A.

**Figure 6.**
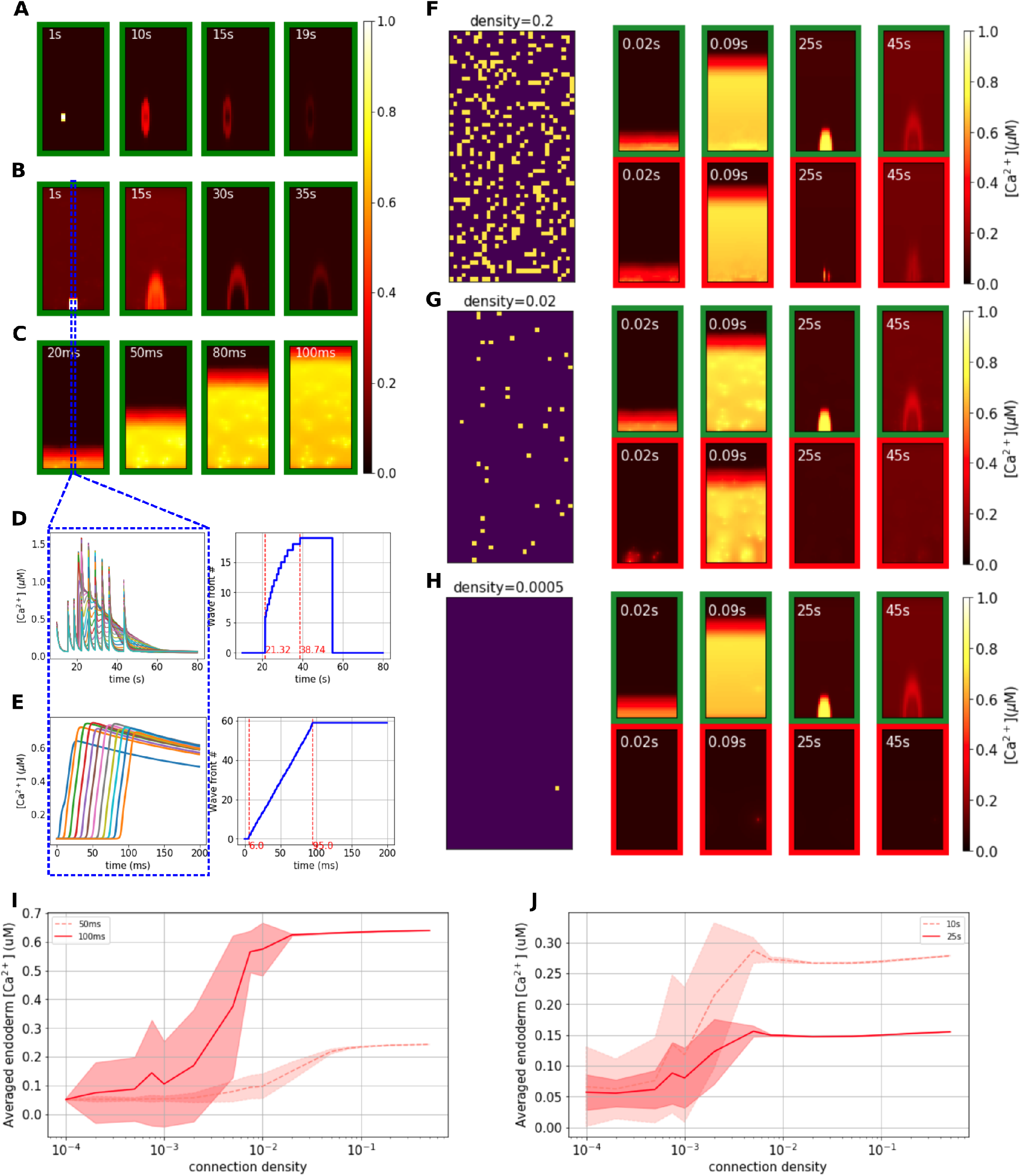
Modeled dynamics of muscle sheets. *(A-C)* Simulation of dynamics under different stimulation patterns in a single muscle sheet, including body column waves *(A)*, bending wave *(B)* and fast wave *(C)*. *(D-E)* [Ca^2+^]_*i*_ traces and wavefronts along the center longitudinal line (marked in the dashed rectangle), shown for the bending wave *(D)* and fast wave *(E).* The right sides show how the wave fronts evolve with time. *(F-H) Connectivity patterns and calcium patterns of ectoderm (green border) and endoderm (red border) in simulations with different connectivity ratios, where the ratios are separately 20% (F), 2% (G) and 0.05% (H). In connectivity patterns each bright spot represents a gap-junctional connection at that position. (I-J)* Diagrams show cross-layer propagation for different connection densities of cross-layer gap junctions. We simulated 20 epochs for each density; we take snapshots at 50ms and 100ms after triggering the fast pathway *(I)*, and 10s and 25s after triggering the slow pathway *(J)*, then average the [Ca^2+^]_*i*_ of the whole endoderm, plotting the mean and standard deviation of the 20 epochs for each density at these time points.

#### Bending wave

A second type of slow wave initiates at the peduncular ectoderm and slowly propagates in the oral direction ***(Szymanski and Yuste, 2019)***. This asymmetrical calcium activity in the ectoderm is believed to cause bending. To simulate the bending wave, we stimulate the slow pathway in a 4×4 cell patch located in the peduncle of the ectoderm sheet of our model; propagation is due as above to gap junctional IP_3_ diffusion, ***Figure 6***B. To obtain a wave as observed in data, the model assumes that gap junctional coupling is anisotropic, larger longitudinally than circumferentially (see Methods).

#### Fast wave

*Hydra’s* contraction burst (CB) is driven by the firing of a unique subnetwork of the ectodermal nerve net ***(Dupre and Yuste, 2017)*** that is distributed across the body but particularly concentrated in a ring in the peduncle. Each CB spike causes a global calcium synchronization which activates all muscle cells in both ectoderm and endoderm. Measurements in ***Szymanski and Yuste (2019)*** showed that the contraction pulses initiate in the peduncular epithelium and propagate to the rest of the body column.

We initiate the fast wave by simulating inputs from a ring of neurons in the peduncle onto the connected muscle cells: we stimulate the fast pathway dynamics in the peduncular row of the ectodermal sheet. Elevated membrane potentials propagate rapidly to the remainder of the cells via their electrical coupling, giving a global calcium synchronization, as shown in ***Figure 6***C.

#### Propagation speeds of wavefronts

We compare the speed of the simulated waves with data. We identified the wavefront of the bending wave in our model as shown in ***Figure 6***D, from which the average propagation speed can be calculated as around 0.9 cells/s. The length of our biomechanical model is in the range of 0.6mm (fully contracted) - 1.7mm (elongated), based on which the length of each cell can be calculated as 9.8 *μ*m (contracted) - 28.8*μ*m (elongated), so the propagation speed is in the range of 9 *μm/s* - 26.5 *μ*m/s. In ***Szymanski and Yuste (2019)***, the propagation speed of the bending wave was calibrated as 13 ± 0.7 *μ*m/s, which falls in the range of the simulation.

The speed of the fast wavefront is computed as shown in ***Figure 6***E, giving 0.7 cells/ms, or 6 mm/s when the model is in the contracted state. This is of the same order as the propagation speed measured from calcium imaging (4.6-5 mm/s) in ***Szymanski and Yuste (2019)***.

#### Cross-layer coupling

Calcium imaging shows that during a contraction burst, neuronal activity is confined to the ectoderm ***(Dupre and Yuste, 2017)***, while both the ectoderm and the endoderm show an almost synchronous fast wave of epitheliomuscular calcium ***(Szymanski and Yuste, 2019**), **Figure 7***. However, body column waves and bending waves are only observed in the ectoderm. The synchronization of fast waves in the two layers is likely due to electrical coupling through the cross-layer gap junctions which penetrate the mesoglea. When cells in the ectoderm are electrically activated, the action potential can propagate through the gap junction and trigger activity in the endodermal epithelium. However, even though cross-layer gap junctions can also allow a flow of IP_3_, there is no evidence that slow waves in ectoderm propagate to the endoderm. We hypothesize that the ability of cross-layer gap junctions to transmit global fast waves but block local slow waves is due simply to their density. We tested this by varying the connectivity ratio of the cross-layer gap junctions in our model. We found when the connectivity ratio is high, both fast and slow waves can propagate between the two layers; when the ratio is low neither of the two waves propagate; and there is a intermediate range in which only the fast wave crosses the layer (***Figure 6***F-H).

**Figure 7.**
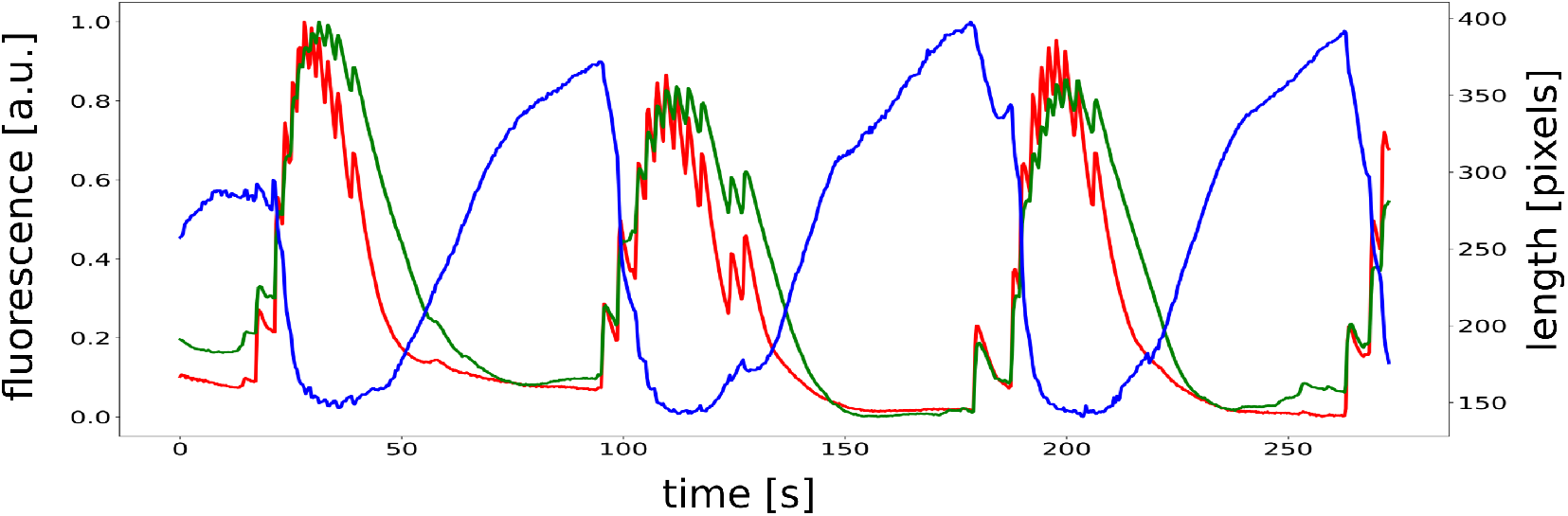
Recorded fluorescence from calcium imaging showing simultaneous activation of ectoderm (green) and endoderm (red) during contractions, shown by computing the corresponding time-varying length of *Hydra.*

To probe the effect of the density of cross-layer gap junctions on the propagation of trans-mesoglea calcium waves, we quantitatively analyzed the relationship between the averaged calcium concentration of the endodermal muscle sheet and the connection density, after stimulating the ectodermal muscle cells’ fast (***Figure 6***I) and slow (***Figure 6***J) pathway. As shown, for both slow wave and fast waves, a higher connection density of cross-layer gap junctions allows stronger calcium wave propagation from the stimulated ectoderm to the endoderm. When the connection density is very low or very high, the standard deviation is small. In intermediate ranges, there is considerable variability, indicating that the specific placement of cross-layer connections can also affect trans-mesoglea propagation. Thus, when the cross-layer gap junctions are distributed under a range of connection density in specific patterns, only the fast waves can cross layers and the slow waves are limited to the directly stimulated layer.

### From neural activity to behaviors

To simulate observed behaviors, we use different spatiotemporal neural firing patterns to drive muscle activation.

#### Contraction bursts

We assume that CBs are initiated by repeated stimulation of the fast pathway in a ring of muscle cells at the peduncle. What causes the CB nerve net to fire is still not known, although it is influenced by conditions such as osmolarity ***(Yamamoto and Yuste, 2019)***, microbes ***(Murillo-Rincon et al., 2017)*** and temperature ***(Tzouanas et al., 2019)***. While the CB nerve net is distributed, it has a high density of neurons at the peduncle and we assume these to be the primarysource of activation. As already shown, gap junctional coupling can propagate this excitation rapidly from the peduncular part to the global body column, activating the whole body. Our choice of cross-layer coupling results in synchronous activation of both body layers, as is observed experimentally ***Figure 7***.

#### Elongation

In principle, the two active muscle layers work against one another: longitudinal contraction by the ectoderm should be countered by the squeezing effect of endodermal contraction creating pressure on the enclosed fluid and stretching the body. If we assume that both muscle layers have the same properties, these active forces indeed counteract one another and our biomechanical model fails to contract (not shown).

It is possible to resolve this paradox if we assume that the two muscle layers have different properties. First, we assume that the same [Ca^2+^]_*i*_. in ectoderm leads to larger active stress than in the endoderm;this is consistent with the fact that ectodermal myonemes are longer than endodermal myonemes ***(Leclère and Röttinger, 2017)***. In this case, longitudinal contraction will dominate. We further propose that ectodermal contraction is *phasic* while endodermal contraction is *tonic*, as a result of a longer-lived latch state, captured by attached dephosphorylated (AM) and attached phosphorylated (AMp) cross bridges in the Hai-Murphy model ***(Hai and Murphy, 1988)***, see Methods. As a result, once neural stimulation ceases, [Ca^2+^]_*i*_. in both layers drops, but the endoderm continues to exert strain. Thus the sustained circumferential contraction gradually exceeds the decaying longitudinal contraction, causing *Hydra* to slowly elongate, in the absence of neural firingor elevated endodermal [Ca^2+^]_i_, as observed experimentally, Fig.***Figure 7***.

#### Bending

*Hydra* frequently initiates bending during contraction. We simulate bending by stimulating the slow pathways of a small localized group of ectodermal cells at the peduncle, which triggers the bending wave as shown in the last section. This generates local contraction which causes *Hydra* to bend towards the stimulated side, ***(Figure 9)***. This shows that activation of the slow and fast pathways can coexist and that the contraction event in the fast pathway does not saturate the calcium dynamics.

#### Nodding

A separate nerve net, called the sub-tentacular network ***(Dupre and Yuste, 2017)***, is found to be correlated with nodding behavior. Here we assume that this network simply stimulates the slow pathways of a small set of ectodermal cells in the sub-hypostomal region. This is similar to bending behavior but with an opposite location. Our model is thus able to generate a “bending” of the hypostome towards the stimulated side ***(Figure 9)***.

Combining these components, we can now simulate complete naturalistic behaviors. Left undisturbed, *Hydra* undergoes repeated cycles of contraction and elongation, combined with bending. We extract the integrated GCaMP fluorescence trace from a neuronal imaging video and use it to infer the firing times of the CB neurons (***Figure 8***A). We further add sparse triggering times for the bending waves (***Figure 8***B). We use this stimulation to drive fast calcium waves and slow bending waves in the epithelial sheets (***Figure 8***D), of which the fluorescence matches the observed curves in ***Figure 7*** (***Figure 8**C*). After encoding the calcium concentration into force (***Figure 8***E, F), we use it to drive the biomechanical model, successfully exhibiting a series of behaviors that mimic the real *Hydra* (***Figure 8***H). The simulated length changes show very good quantitative agreement with the dynamics of contraction and elongation of the animal (***Figure 8***G).

**Figure 8.**
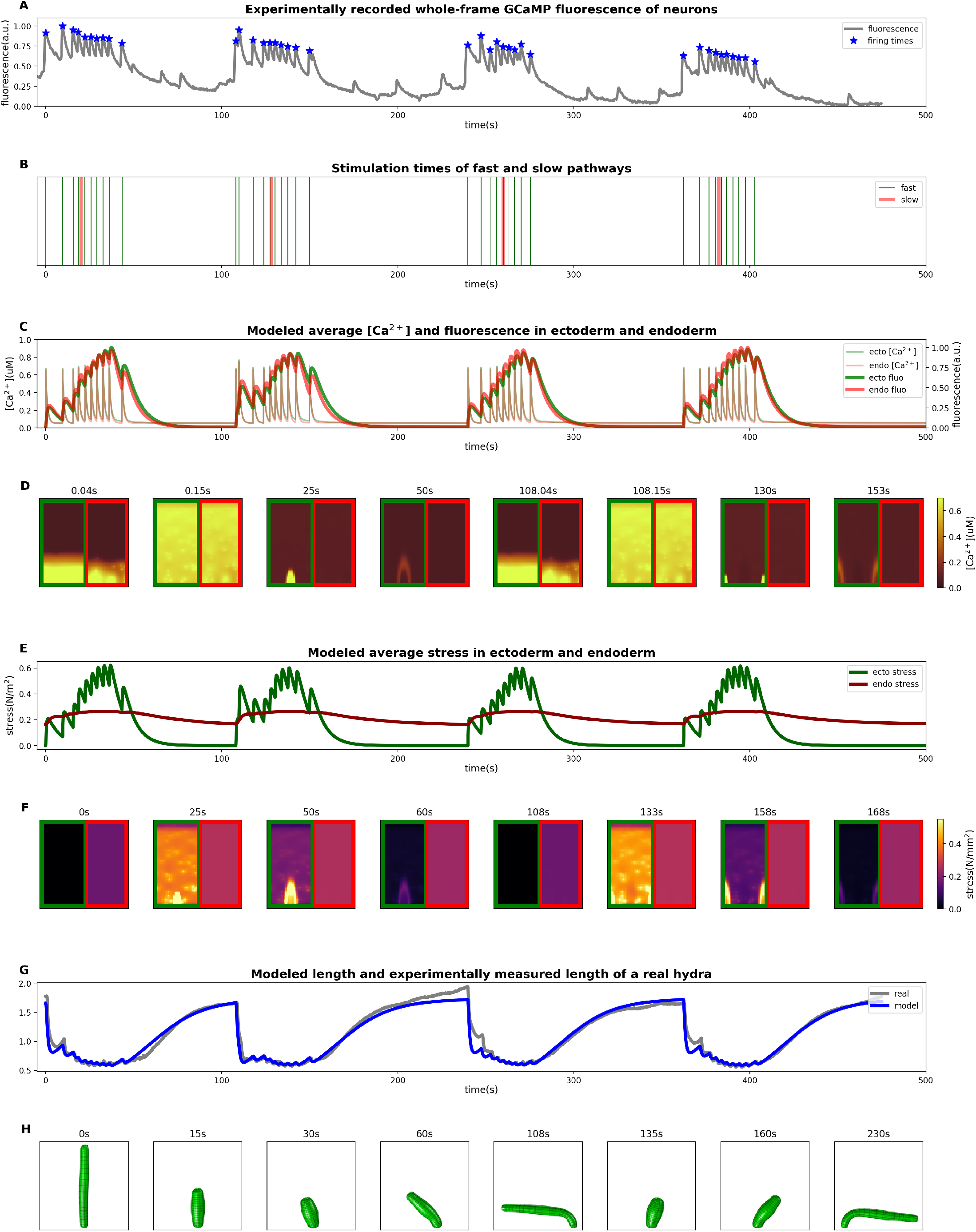
Simulation from stimulation to the behaviors. *(A-B)* NGCaMP fluorescence trace: stars mark estimated times of neural firing *(A)* and consequently, stimulation times for the model *(B)*. *(C-D)* Averaged [Ca^2+^]_*i*_ and fluorescence intensities in ectoderm and endoderm *(C)* and [Ca^2+^]_*i*_ patterns of some moments *(D)*. *(E-F)* Averaged active stress in ectoderm and endoderm *(E)* and stress patterns of some moments *(F)*. *(G-H)* Comparison between length evolution of the model and from a *Hydra (G)* recording and some stills of the final simulated behaviors from the model *(H)*.

With designed or measured neural stimulation as input, our model successfully simulated the behaviors of contraction, bending and elongation as shown in ***Figure 9***.

**Figure 9.**
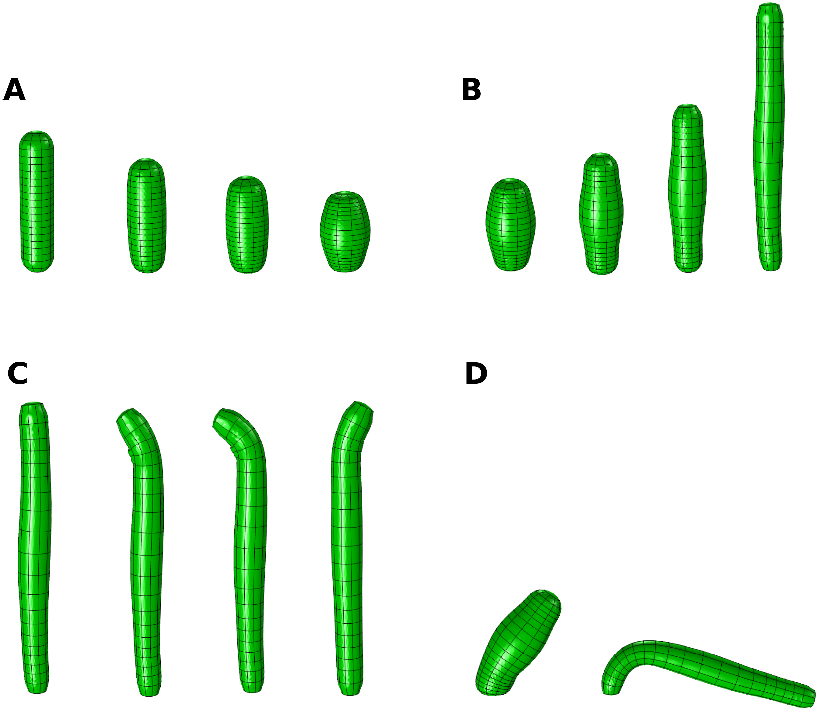
Different behaviors simulated by the model. *(A)* contraction *(B)* elongation *(C)* nodding *(D)* bending.

## Discussion

In this work, we have succeeded in implementing a model framework that predicts behaviors from neural firing patterns. Here we review the assumptions, required mechanisms and limitations of our model. We began with a biophysical model for single muscle cell dynamics that is grounded in known molecular mechanisms;we then linked muscle cells into a network via gap junctional coupling, which successfully simulated multiple-timescale calcium activation dynamics observed in GCaMP imaging experiments in *Hydra.* Such multiple-timescale calcium signaling was also recorded in arterial smooth muscle cells ***(Halidi et al., 2011)***. We then used this model to generate active stress to drive a passive biomechanical model of the body. This model successfully simulated *Hydra* behaviors including contraction bursts, bending and elongation. These two components together form a model that can exhibit different behaviors with given neuronal stimulation, and can thus serve as a testbed for reverse engineering neural activity needed for each behavior of *Hydra.*

In order to match observed activity and behavior, we raised and addressed a series of questions. We accounted for the two distinct time scales of calcium patterns in imaging experiments by assuming that there are two different calcium pathways (ionotropic and metabotropic) in each single muscle cell, then captured the different propagation speeds through the dual functions (electrical coupling and chemical diffusion) of gap junctions. By assuming sparsely distributed cross-layer gap junctions between ectoderm and endoderm, we succeeded in producing synchronized fast waves in the two layers along with the isolation of slow waves to the ectoderm. To explain how *Hydra* can longitudinally contract although the two counteraligned muscle layers are simultaneously activated, we postulated that muscles of ectoderm and endoderm have different properties (phasic and tonic). This further explains how *Hydra* can elongate with no apparent endodermal calcium activation ***(Figure 7)***. Putting these factors together, we successfully simulated cycles of contraction, elongation and bending.

Despite this success, there are many details and limitations that need to be explored further. For instance, how and which groups of neurons transmit these distinct stimuli to the muscle is still unclear. We have considered a simplified structure of *Hydra*, and neglected the effects of the surrounding water, whose viscosity and buoyancy may influence behavior ***(Naik et al., 2020; Megill, 2002; Rudolf and Mould, 2009)***. Furthermore, our model only assumes feedforward transformations from neural activity to behaviors, although it is likely important to understand how behavior influences neural firing through mechanosensation and other forms of sensory feedback.

### Neuromuscular transmission in *Hydra*

An open question is how and where neurons connect to and stimulate muscles. Electron microscopic observations showed clear evidence of neuromuscular synaptic junctions in *Hydra* tissue ***(Westfall, 1973)***. Further studies have shown that the neuropeptides Hydra-RFamides and Hydra-KVamide (expressed in the peduncle region) may play roles in neuromuscular transmission ***(Yum et al., 1998; Hansen et al., 2000; Gründer and Assmann, 2015)***. The neuropeptide Hym-176C can induce ectoderm contraction and is selectively expressed in ectodermal peduncle neurons ***(Yum et al., 1998; Siebert et al., 2019; Noro et al., 2019; Klimovich et al., 2020)***. Non-selective cation channels HyNaCs were identified in epitheliomuscular cells, which are directly activated by Hydra-RFamides I and II and can depolarize the cellular membrane potential ***(Dürrnagel et al., 2012; Gründer and Assmann, 2015)***. In our model, we propose that two different types of neuropeptides play the roles of triggering different pathways (see Methods), enabling differential control of different timescales dynamics in the muscle layers, potentially by distinct neurons.

### Electrical signaling in nerve net and muscle

Our model treats the question of whether electrical activity during contraction bursts is transmitted through the nerve net or the muscle layer. We propose that hypostomal neurons play a role of integrating information from the environment: following a decision to fire, the signal is propagated through the sparsely distributed CB nerve subnet to the peduncle ring of motor neurons, which acts as the primary drive of contraction. This architecture is supported by the observed propagation of fast calcium activation from the peduncle towards oral side ***(Szymanski and Yuste, 2019)***; the expression of Hym-176C in the peduncle (***Yum et al., 1998)*** and the previous observation of electrical conduction in the nerve-free *Hydra* epithelia ***(Campbell et al., 1976)***.

However, we believe that during contraction, synchronous drive from the nerve net is supplemented by the electrical coupling property of muscle cells themselves. The CB neurons are sparsely distributed in the body column; it is unlikely that the CB neurons directly innervate all muscle cells, although it is possible that each neuron drives a small group of them. We simulated the dynamics that result from setting the electrical conductance between muscles to zero and stimulating random small groups of muscle cells; this leads to slowly growing nodes of excitation via the slow chemical diffusion of IP_3_ ***(Figure 10)***. Thus we believe that electrical coupling of muscle cells is needed to explain the rapid synchronization of calcium activity in the epithelium ***(Badhiwala et al., 2020)***. The distributed network of CB neurons are likely necessary to integrate and generate the contraction activity, and may contribute to the robustness of contraction.

**Figure 10.**
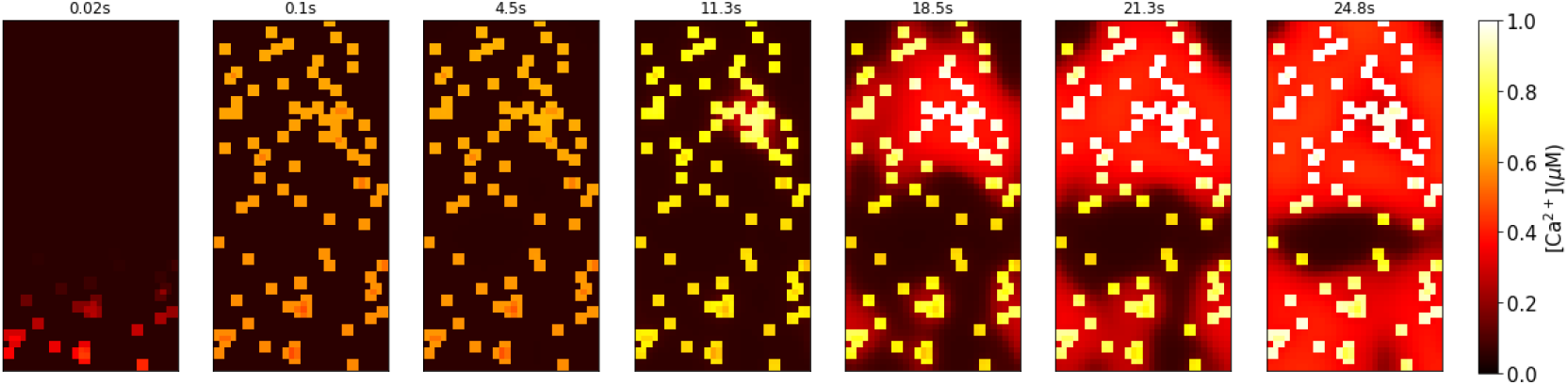
Modeled calcium activation pattern under sequential stimulation of the fast pathways of randomly chosen 2 by 2 groups of muscle cells in the body column with no gap junctional electrical conductance.

Here we demonstrated that sparse electrical coupling between the muscle layers can account for the coactivation of longitudinal and circumferential contraction during contraction bursts. While it is counterintuitive that these muscles would activate together, this may have a functional role; as suggested in ***Szymanski and Yuste (2019)***, the resulting stresses on the body may serve to squeeze absorbed water out of the body walls.

*Hydra* has two additional known synchronously firing networks, the “rhythmic potentials” RP1 and RP2. We have found no correlation of RP firing with instantaneous length changes; thus it is unlikely that the RP networks directly control muscle contraction. However, it has been suggested the “radial contraction” behavior is particularly related to RP2 ***(Dupre and Yuste, 2017)***. The ectodermal RP1 nerve net may inhibit CB neurons therefore suppressing contraction bursts, based on the fact that its frequency is inversely proportional to contraction bursts ***(Passano and McCullough, 1962; McCullough, 1965; Dupre and Yuste, 2017)***.

### Feedback from behavior and the environment

Our model simulates the transformation from neural stimulation to behaviors as a purely feedforward control pipeline, but there are several ways in which feedback may play a role. The rules that govern the generation of the neural activity are still an open question. Recent experiments have shown that temperature ***(Tzouanas et al., 2019)*** and osmolarity ***(Yamamoto and Yuste, 2019)*** can influence the CB firing rate in *Hydra.* Neural activity is likely to be influenced by the behavior or the state of the animal through mechanosensation and other sensory inputs. This may occur through direct sensory feedback or through alternate mechanisms such as changing ionic concentrations in the intercellular medium, the external solution or the interon.

A further source of potential feedback that we neglect is that between the movement of the animal and the dynamics of diffusion. In the model, we assumed that the speed of IP_3_ diffusion is faster in the longitudinal direction than in the circular direction (see Methods), in order to account for the different propagation speeds of bending waves in longitudinal and circular directions, in units of cells per second. While it is possible that the density of longitudinally oriented gap junctions may be larger than that of circularly oriented ones, it is also possible that the propagation speeds in the two directions are the same, but since bending waves are usually initiated when *Hydra* is contracted and cells are squeezed longitudinally, the wave may travel through more cells longitudinally than circumferentially. In order to incorporate this, one would need to model the relationship between the coefficient of IP_3_ diffusion and the local cell shape. Implementing such feedback between the biophysics of the muscle layer and the geometry of the biomechanical model would require considerable engineering effort. In general, however, accounting for any of these effects is possible by extending the framework of our model.

### Body plan simplicity

Our biomechanical model simplifies *Hydra* to a hollow cylinder with two spherical ends. However, *Hydra’s* body column deviates from cylindrical, slimming toward the peduncle. Furthermore, cells across the body are heterogenous in size and shape. Recent work indicates that there is variation in the Young’s modulus of the body column, and suggests that this can affect somersaulting behavior ***(Naik et al., 2020)***. Future work could explore more precisely how the body wall mechanics transform neural signals and muscle forces into behavior. Further, we do not attempt to model the tentacles, whose sensory input likely contributes significantly to *Hydra’s* movement, and whose adhesion to surfaces frequently affects body movement. Modeling such details will be necessary to obtain detailed quantitative agreement between the model and additional aspects of behavior. Our goal here was to capture the most significant factors of the biomechanics that underlie the behaviors of *Hydra* that have to date been recorded simultaneously with calcium imaging, excluding higher order complexities. We hope our model will serve as a starting point for further work to capture the full richness of *Hydra’s* natural behavior.

### Significance

Our study shows that in *Hydra*, with one of the simplest nervous systems, the functions of neurons and muscles are not fully segregated - behavior is due to an interplay of nerve and muscle excitation, transformed by body mechanics. Besides its relevance to the understanding of neuromechanical coupling in excitable tissues like the heart or the gut, our work demonstrates the feasibility of a complete model, from the neuronal basis to muscle and body biomechanics, of an animal behavior.

## Acknowledgments

This project is supported by NSF CRCNS 1822550 and a Simons Collaboration for the Global Brain grant to A.F.. R.Y. was also supported by the Burroughs Wellcome Fund 2018 Collaborative Research Travel Grant and the Vannevar Bush Faculty Award (ONR N000142012828). MBL research was supported in part by competitive fellowship funds from the H. Keffer Hartline, Edward F. MacNichol, Jr. Fellowship Fund, The E. E. Just Endowed Research Fellowship Fund, Lucy B. Lemann Fellowship Fund, Frank R. Lillie Fellowship Fund Fellowship Fund, the Fries Trust Research Award, Hartline MacNichol Research Award, L. & A. Colvin Summer Research Fellowship, and John M. Arnold Fellowship Research Award of the Marine Biological Laboratory in Woods Hole, MA. M.B., A.F. and J.S. were supported by the University of Washington’s Royalty Research Fund. We thank Rob Steele, Celina Juliano and Jacob Robinson for valuable discussions. We thank the MBL Whitman Center for supporting the Hydra lab in the summers of 2017, 2018 and 2019.

## Notes

### Competing Interest Statement

The authors have declared no competing interest.

